# Application of UV-cured resin as embedding/mounting media for practical, time-saving otolith specimen preparation

**DOI:** 10.1101/474643

**Authors:** Carlos Augusto Strüssmann, Kaho Miyoshi, Shota Mitsui

**Author notes:** Corresponding author: Carlos Augusto Strüssmann, Graduate School of Marine Science and Technology, Tokyo University of Marine Science and Technology, 4-5-7 Konan, Minato-ku, Tokyo 108-8477, Japan. **E-mail address of co-authors**: Kaho Miyoshi Shota Mitsui.

## Abstract

Otoliths are calcified structures located in the inner ears of fish, as in most vertebrates, that are responsible primarily for the perception of gravity, balance and movement, and secondarily of sound detection. Microstructural and chemical analyzes of the inner otolith growth layers, called increments, constitute powerful tools to estimate fish age and elucidate many life history and demographic traits of fish populations. Otolith analyzes often require the production of a thin cross section that includes in the same plane of view the otolith core and all microscopic layers formed from birth until the moment of collection (otolith edge). Here we report on the usefulness of UV-cured resins that have become recently popular among nail artists and hobbyists for otolith specimen preparation. We show that single-component UV-cured resins can replace successfully and advantageously the commonly used two-component Epoxy resins to obtain otolith cross sections suitable for both microstructural examination and chemical analysis by electron probe microanalysis. UV-cured resins provide on-demand, extremely rapid (minute-order) hardening and high transparency, while providing similar adhesion and mechanical support for the otoliths during processing and analysis as Epoxy resins. UV-cured resins may revolutionize otolith specimen preparation practically- and time-wise, and may be particularly useful in teaching and workshop situations in which time for otolith embedding is a constraint.

## 1 Introduction: otoliths in fisheries science and resource management

Otoliths are calcified structures located in the inner ears of fish, as in most vertebrates, that are responsible primarily for the perception of gravity, balance and movement, and secondarily of sound detection. Some invertebrates like cephalopods also possess similar structures called statoliths (Dilly, 1976). Fish otoliths grow in size as fish age by consecutive accretion of layers, called increments, in proportion to somatic growth under a species-specific and environment-dependent relation (Stormer and Juanes, 2016). The increments are composed mainly of crystalline calcium carbonate embedded in an organic matrix and are layered consecutively on the outer surface of the otolith starting from a microscopic core formed during early embryonic development (Campana and Nielson, 1985; Watanabe and Kuji, 1991; Morales-Nin et al., 2005). Calcium carbonate and organic matter deposition follow primarily a diel rhythm, with the former predominating at daytime and the latter at nighttime, and secondarily a seasonal rhythm that alternates periods of intense and reduced somatic growth (Mugiya, 1987; Kono et al., 2014).

Analyses of the otolith microstructure and chemistry constitute powerful tools to elucidate life history and demographic traits of fish populations (Stevenson and Campana, 1992; Campana et al., 2016; Neville et al., 2018). The incremental, patterned deposition of materials on the otolith surface can be visually recognized as “annual” and “daily” rings whose identification provides information on an individual’s age in years or even in days and therefore can be used to clarify its year or even date of birth (Tsukamoto et al., 1989; Fowler, 1990). Likewise, the width of the daily and annual increments provides information on the amount of somatic growth during a particular day or period in life (Jones, 1992; Castellini et al., 2017; Watai et al., 2017). On the other hand, the chemical composition of the daily increments is affected by the physiological status of the fish and by the surrounding habitat’s abiotic conditions such as water chemistry and temperature (Radtke, 1989; Campana, 1999; Secor and Rooker, 2000). Unlike in true bones whose constituents can be resorbed under periods of food deprivation or environmental stress, the chemical composition of the otolith increments is fixed for life (Campana, 1983; Ichii and Mugiya, 1983). Thus, chemical analysis of individual increments, which as noted earlier can be ascribed to particular ages or life stages, may provide critical information on the current and past environmental experiences and physiology-altering events (e.g. reproduction, metamorphosis, migrations, settlement, etc) of individual fish (Zenitani et al., 2007; Hamilton and Warner, 2009; Shiao et al., 2010; Grønkjær, 2016; Arai and Chino, 2018).

## 2 Current methods for otolith specimen preparation

The methodology on fish otolith specimen preparation has been elegantly reviewed in Stevenson and Campana (1992). Small otoliths such as those from larval and juvenile stages often are thin and transparent enough to allow visualization of the increments by simple clearing and mounting on a glass slide. However, structural analysis of otoliths from medium to large-sized specimens and chemical analysis of the inner increments in otoliths of any size require the production of a more or less thin cross section of the otolith that exposes in the same plane of view the otolith core and all layers formed from birth until the moment of collection (otolith edge). Otoliths are hard but relatively brittle and may be small and difficult to process into a section without support. Thus, cross sections are usually obtained after embedding it to produce a block or mounting it on a glass slide or prop with a liquid media that is subsequently hardened (cured) by physical (e.g. heat) or chemical (e.g. catalyzer) means and which would support the otoliths during the sectioning/lapidating/polishing process. The techniques and media used for otolith embedding or mounting have been reviewed by Secor et al. (1992). Otolith researchers have experimentally used a variety of resins, glues, or waxes in otolith specimen preparations but by far the majority of the otolith studies published used some variant of Epoxy resins. This is probably due to their high transparency and chemical stability during long term storage.

Each type of medium used for otolith specimen preparation have advantages and disadvantages and very often these tradeoffs limit their applicability. Embedding media properties include final hardness, which affects the easiness of cutting and polishing, permeability into or adhesion to the surface of the otoliths (associated with differences in medium viscosity and hydrophilicity), transparency and/or occurrence of air bubbles before or after hardening, which affects observation by transmitted light microscopy, chemical stability during storage or chemical analysis (for example, while under an electron beam for electron probe microanalysis), and many others. One important property is the time for hardening, which can be troublesome both if the embedding medium starts hardening too soon when the otolith is still being oriented in the molding cast or too slow, meaning that several hours to days will be spent until a casting of sufficient hardness for processing can be obtained. It is beyond the scope of this report to compare the advantages and disadvantages of all available embedding/mounting media for otoliths, for which the reader is referred to Secor et al. (1992).

## 3 Advantages of UV-cured resins for otolith embedding and specimen properties

Here we report on the usefulness of UV-cured resins that have become recently popular among nail artists, DIY jewelers and other hobbyists, to produce otolith cross sections suitable for both microscopical examination of structure and chemical analysis by electron probe microanalysis (EPMA). We have experienced with the acryl acrylate type of UV-cured resins for the past 18 months and have come to the conclusion that they can replace advantageously other types of resins like the traditionally used Epoxy resins in most, if not all situations. The basic properties of UV-cure resins that relate to otolith preparation are as follows.

1) On-demand, extremely rapid curing. Practical, grinding/polishing-level hardness is obtained within minutes depending on the block thickness and power of the UV light source. In contrast, traditional one- or two-component Epoxy resins may take hours or days to harden depending on temperature and otoliths embedded in them occasionally change their position (shift, twist, turn) during hardening. Of equal importance, hardening only starts under illumination with the appropriate wavelength (in this case c.a. 365–400 nm). This ensures unlimited working time for otolith observation and orientation during embedding/mounting, but conversely, that otoliths can be immobilized in the desired orientation almost immediately by turning on a UV light source (Fig. 1).

**Fig. 1.**
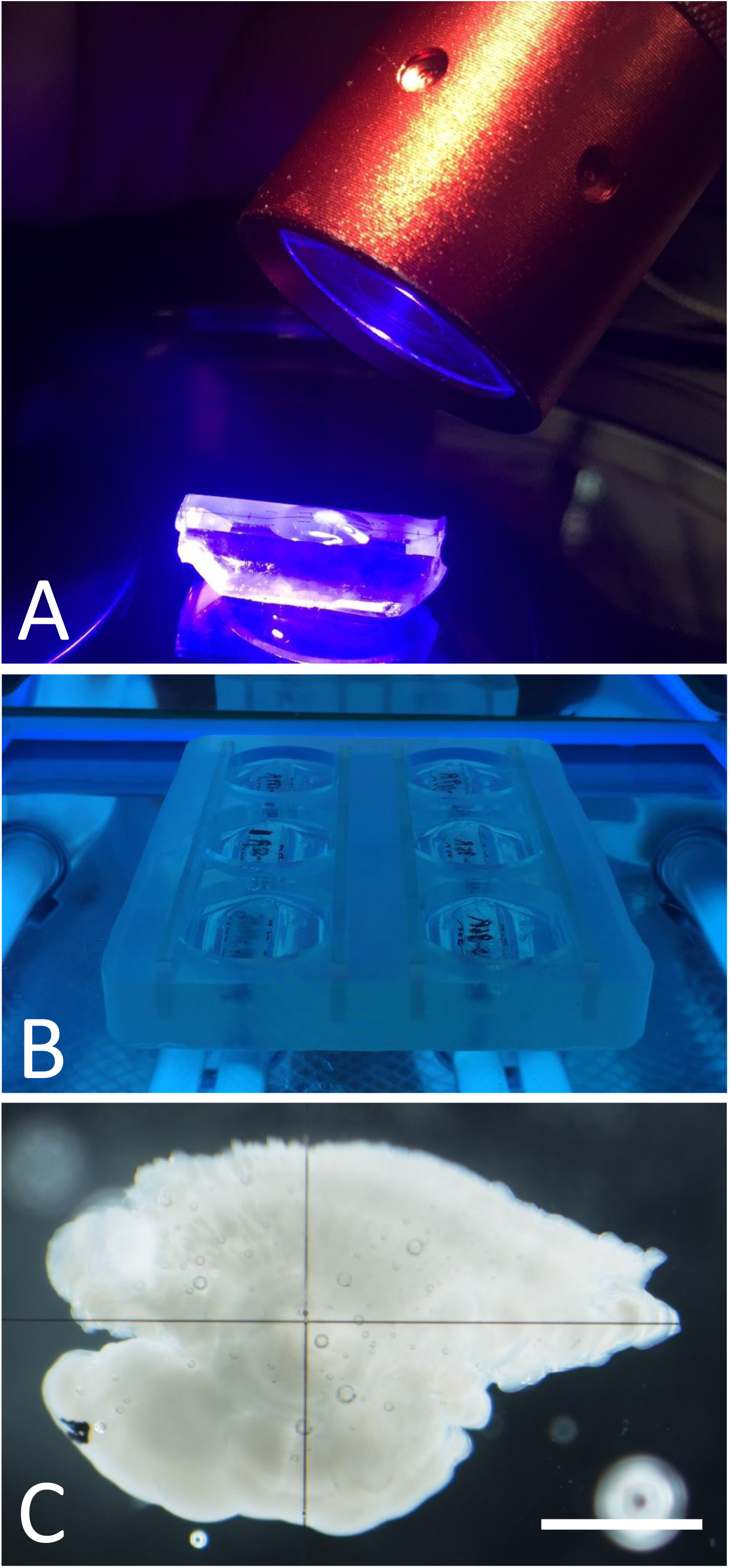
Embedding of otoliths using UV-cured resin. A: Fixation of otolith onto a grinding/polishing guiding block using UV-cured resin; the block is illuminated for about 15 s with a hand-held UV lamp (400 nm) while still on the microscope stage. B: Hardening of 25 mm-diameter round grinding/polishing blocks including the guiding block+otolith produced in A; hardening of the blocks is obtained by illumination for 5-15 min in a commercially available chamber commonly used in nail salons (UV lamp 365 nm; lamp power 9W). C: Appearance of a chum salmon *Oncorhynchus keta* otolith fixed onto a guiding block and subsequently embedded in the center of a block of UV-cured resin with total thickness of 25 mm (one side thickness of 12.5 mm); note the high transparency of the hardened resin. Note also crosshair lines on the surface of the guiding block that are used for otolith axis and nucleus orientation during sectioning/grinding. Scale bar represents 1 mm.
2) High transparency before and after hardening. This ensures that otoliths can be clearly visualized inside the blocks at any time. This property is critical when attempting to locate the position of the otolith core during embedding (before hardening) or during cutting/grinding (after hardening) (Figs. 1, 2). This characteristic is particularly suited for using orientation blocks with guidelines to help locate the approximate position of the nucleus and the distance remaining from the grinding/polishing edge to the nucleus during processing as we perform in our laboratory (Strüssmann CA and Colautti DC; Patent pending).

**Fig. 2.**
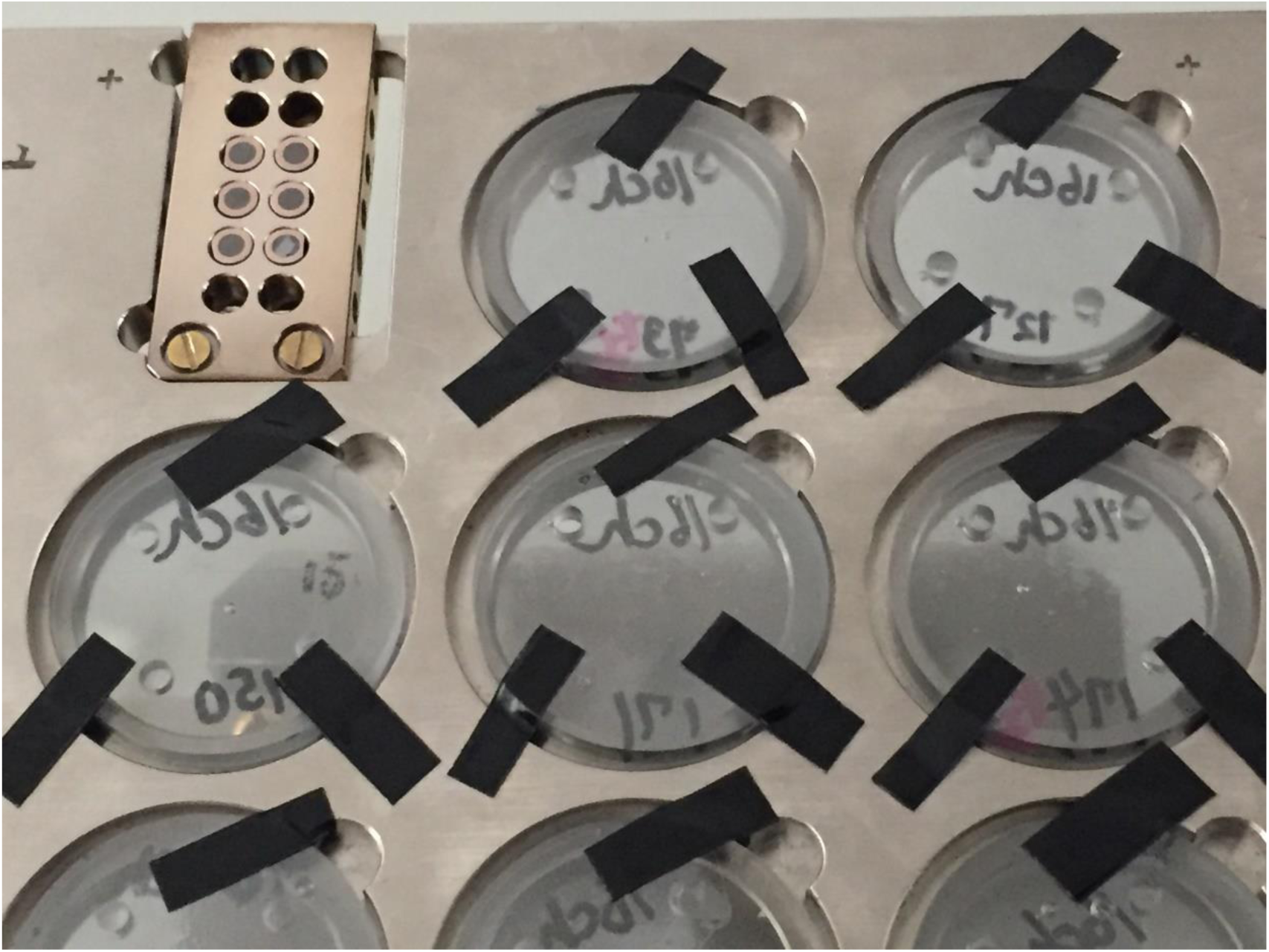
Appearance of round slides (25 mm diameter, 1-1.2 mm thick) made of UV-cured resin and which contain otolith specimen cross sections ready for EPMA analysis. The slides can be mounted directly on the specimen holder of the EPMA equipment. Note the transparency of the UV-resin blocks in spite of being already coated with Pt-Pd for EPMA (or SEM) analysis.
3) Sufficient hardness for grinding and polishing. UV-cured resins provide adequate mechanical support for the otoliths during processing until the obtention of cross sections (Figs. 3–5). This is important to prevent cracking of the otoliths that are common with softer, fast-hardening embedding/mounting media such as thermoplastic glues or waxes. Shrinkage may be slightly higher than with traditionally used Epoxy resins and this could be a problem for small or brittle otoliths, but we have not experienced any significant problems to date.

**Fig. 3.**
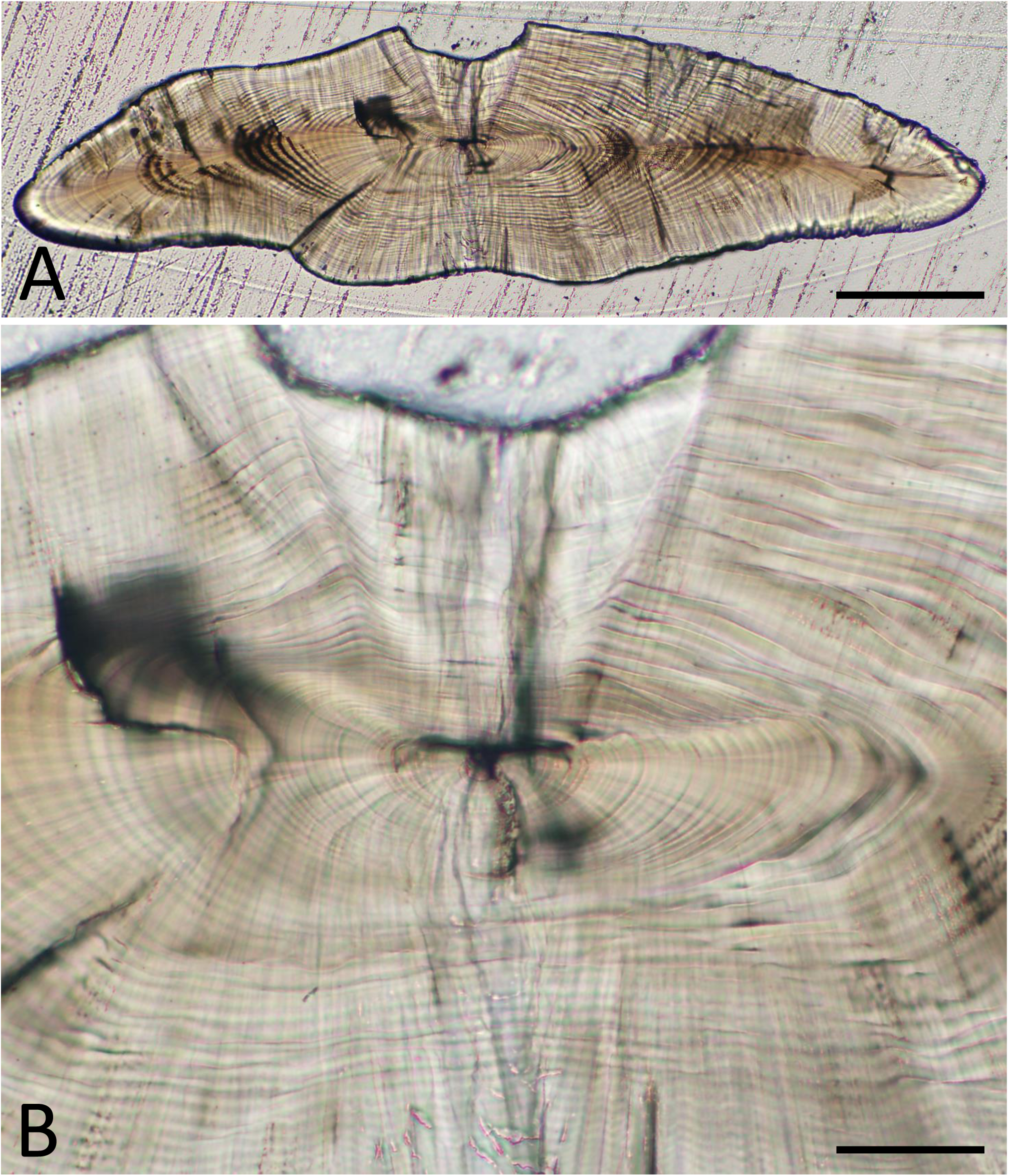
Transverse, thin section of a cobaltcap silverside *Hypoatherina tsurugae* otolith embedded with UV-cured resin. Scale bars represent 200 and 50 μm in A and B, respectively.

**Fig. 4.**
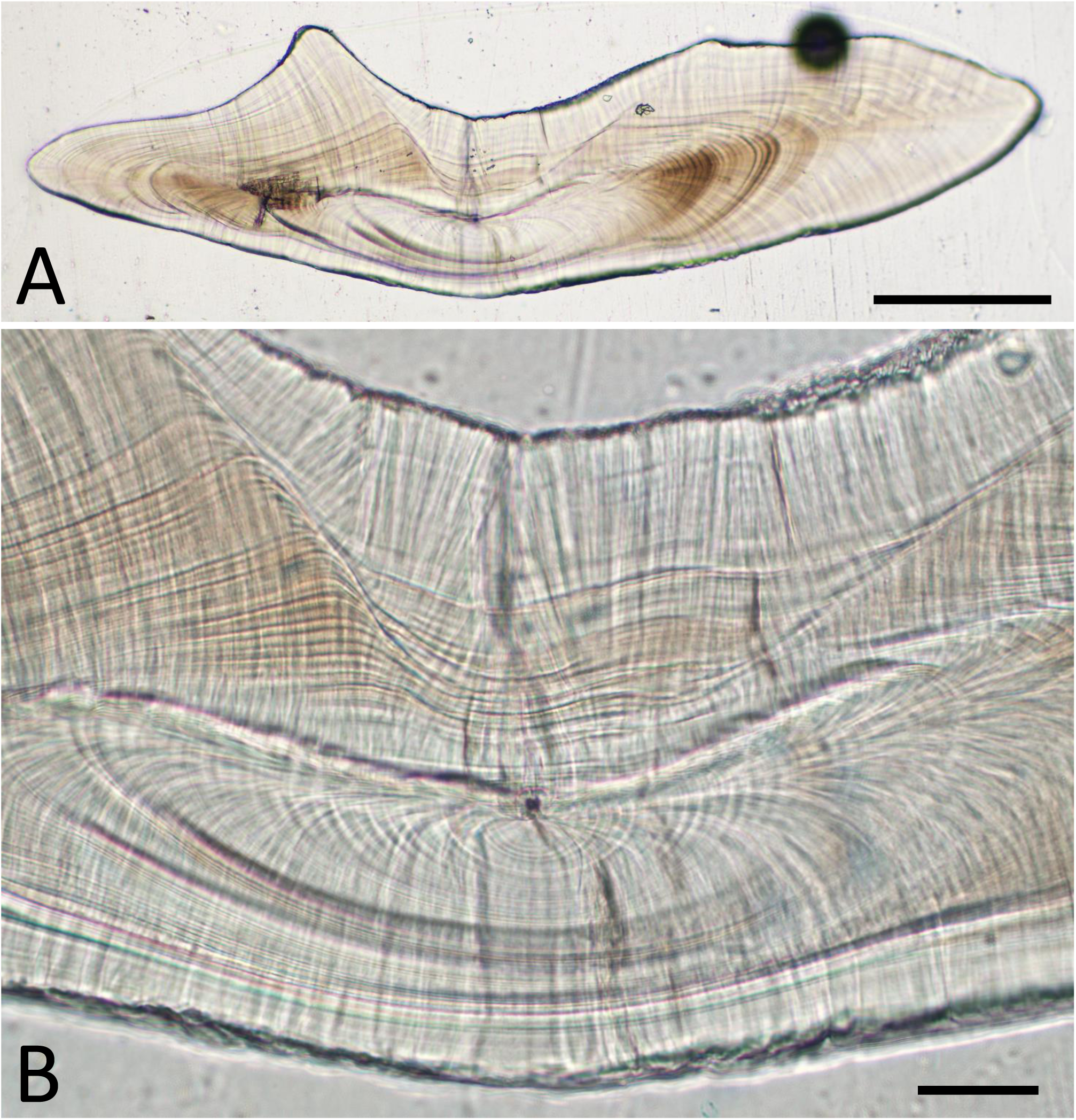
Transverse, thin section of a round herring *Etrumeus teres* otolith embedded with UV-cured resin. Scale bars represent 200 and 50 μm in A and B, respectively.

**Fig. 5.**
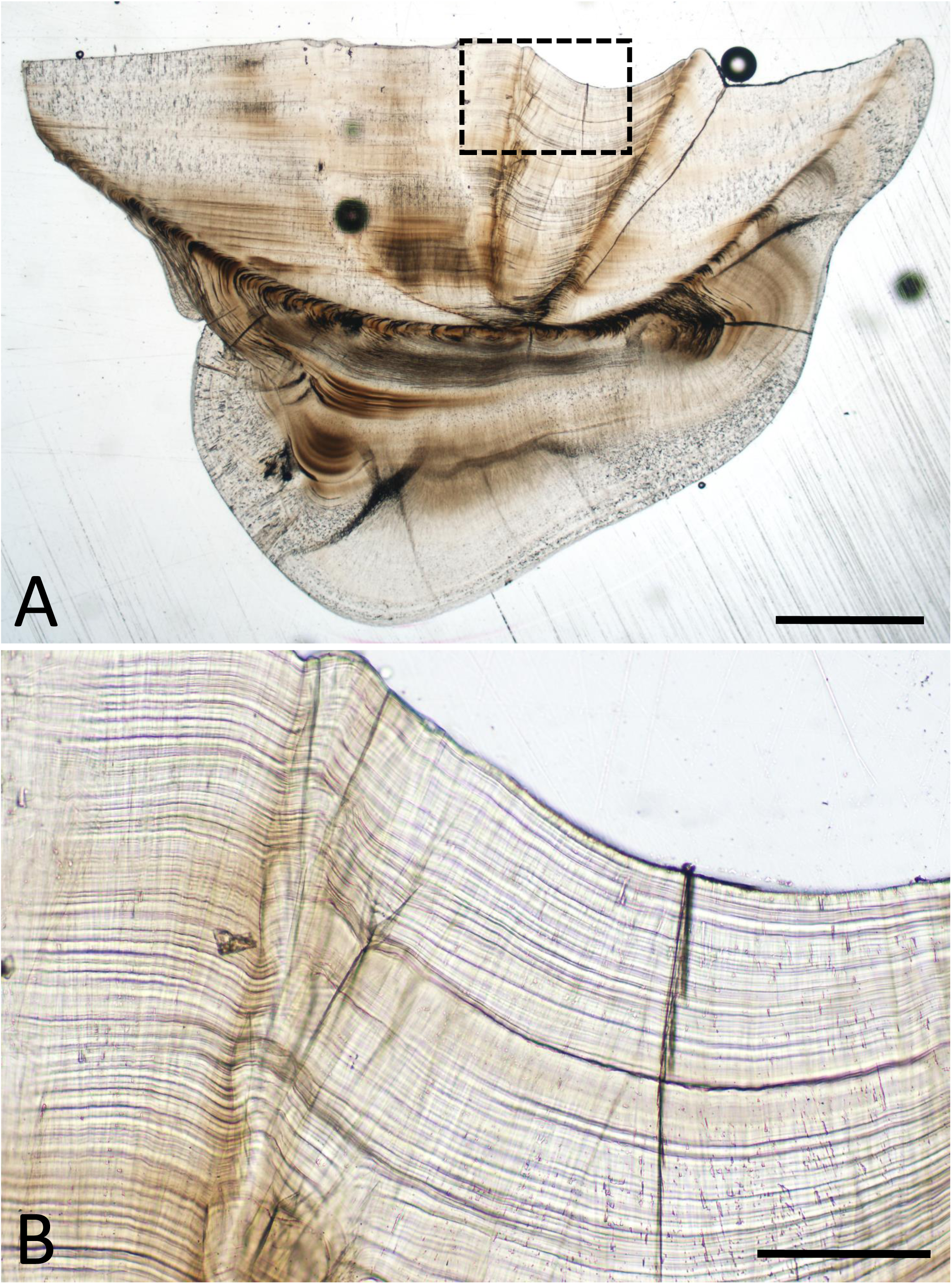
Transverse, thin section of a silver croaker *Pennahia argentata* otolith embedded with UV-cured resin. Scale bars represent 1000 and 200 μm in A and B, respectively.
4) Strong adhesion to the otoliths. The otoliths become firmly attached to the resin and do not detach during wet grinding or polishing. We have not yet tested if adhesion is sufficient for cutting with a precision cutter, so this aspect needs further testing.
5) Thermoplastic stability under an electron beam. UV-cured resins seem to be as stable under an electron beam as the Epoxy resins traditionally used for embedding of EPMA specimens (Fig. 6). The results of semi-quantitative EPMA analysis of UV-cured and Epoxy resins indicate a slightly different chemical composition, e.g. higher Sulphur content of UV-cured resin vs higher Chlorine content for Epoxy resins (Fig. 7) but this has no bearing on the results of otolith chemical composition (see 6). Of great importance, in over 18 months of use for EPMA analyses of hundreds of otolith specimens we not noticed any change in the rate of EPMA column contamination due to the use of UV-cured resin as compared to Epoxy resin.

**Fig. 6.**
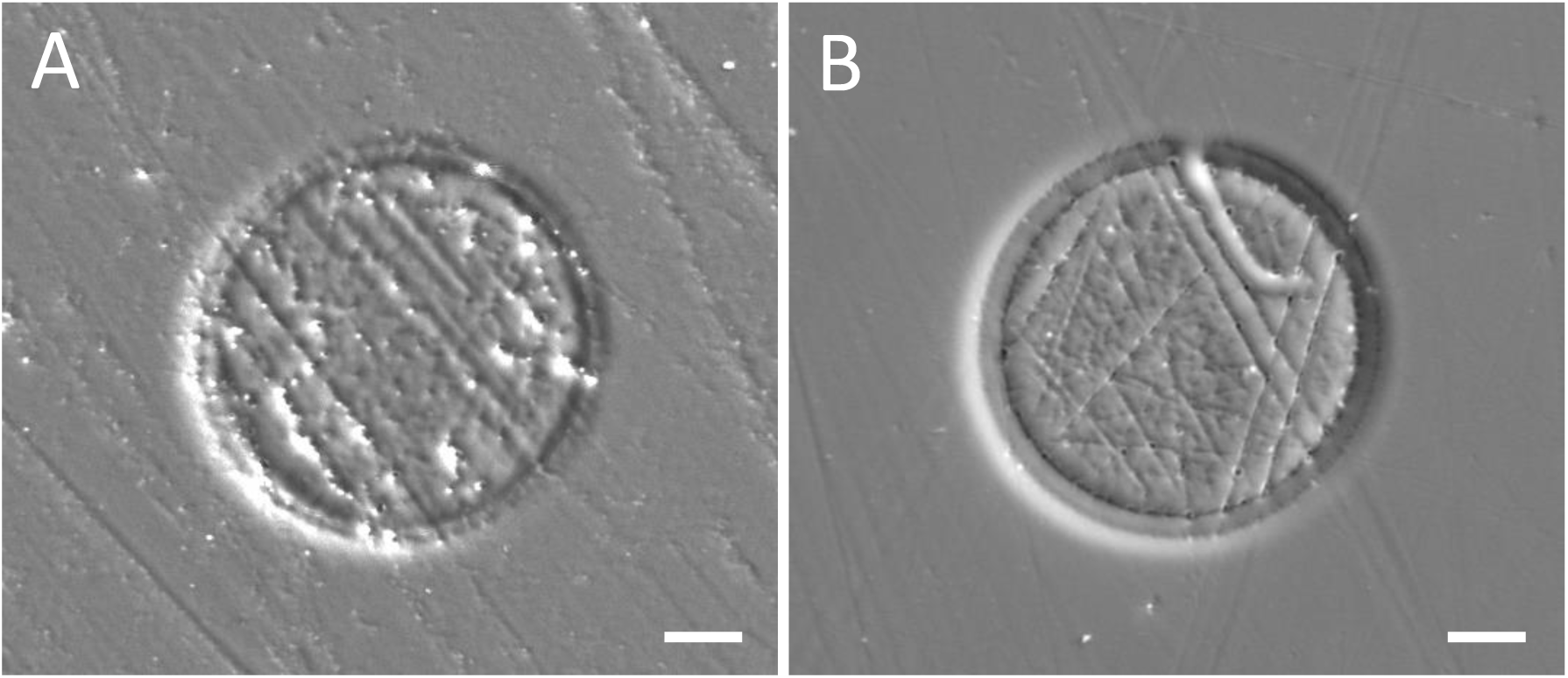
Comparative deformity on the surface of Epoxy and UV-cured resin blocks irradiated with an electron beam (diameter of 50 μm) for 100s during EPMA analysis. The results suggest comparable thermoplastic stability for UV-cured resin and Epoxy resin under an electron beam. Scale bars represent 10 μm.

**Fig. 7.**
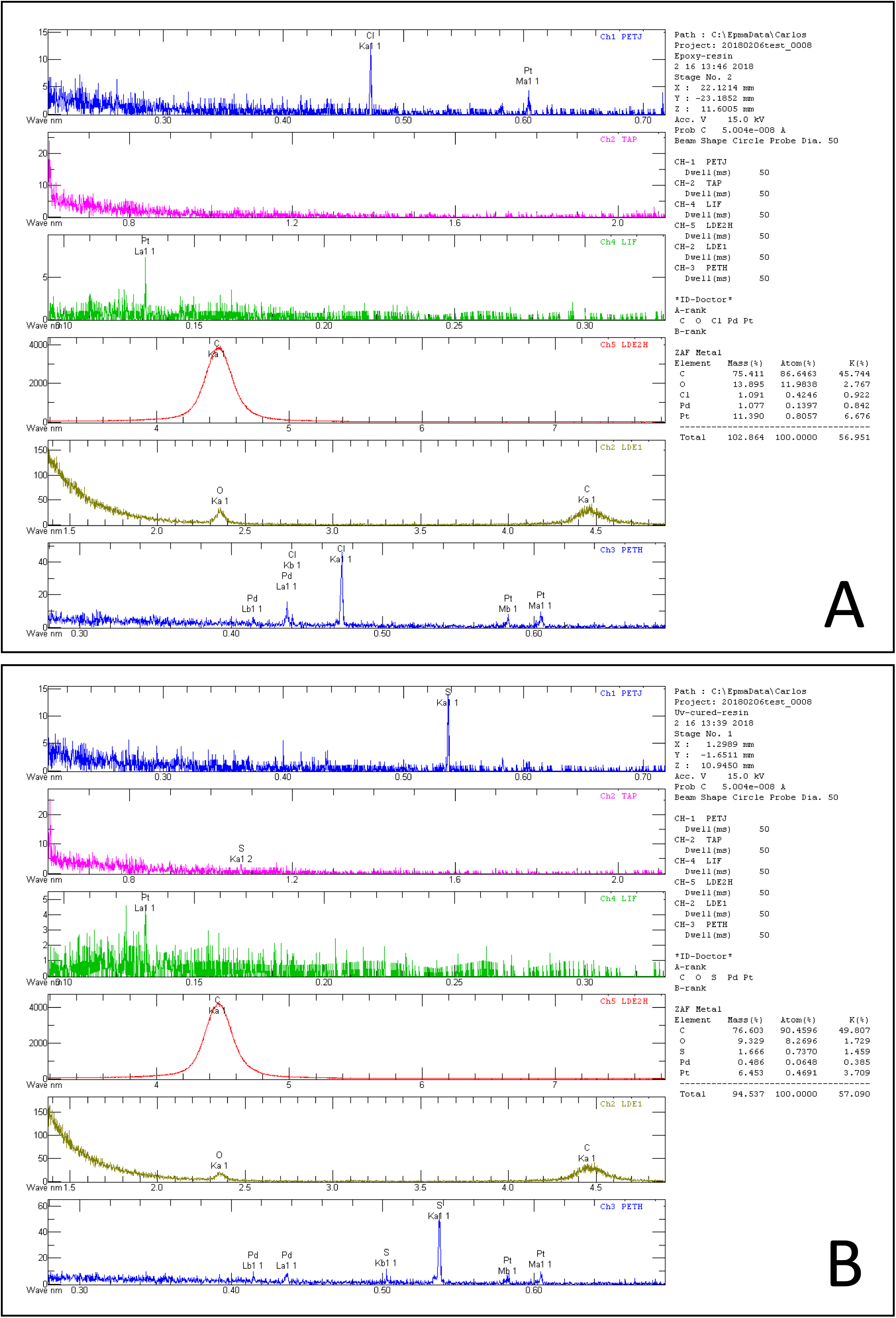
Typical results of semi-quantitative EPMA analysis of Epoxy (A) and UV-cured (B) resin blocks. Results reveal the characteristic presence of Chlorine and Sulphur in Epoxy and UV-cured resins, respectively.
6) Finally, as concerns the chemical analysis of otoliths for reconstruction of the environmental history of individual fish, utilization of UV-cured resins do not affect significantly the elemental composition of otoliths as determined by EPMA (Figure 8). A note of caution is that otoliths embedded in UV-cured resin appear to have somewhat higher Calcium values than those in Epoxy resin. However, this would only be a problem when trying to compare results obtained with specimens prepared by different methods. Moreover, we have not tested the effects of UV-cured resin embedding on the isotopic composition of otoliths.

**Fig. 8.**
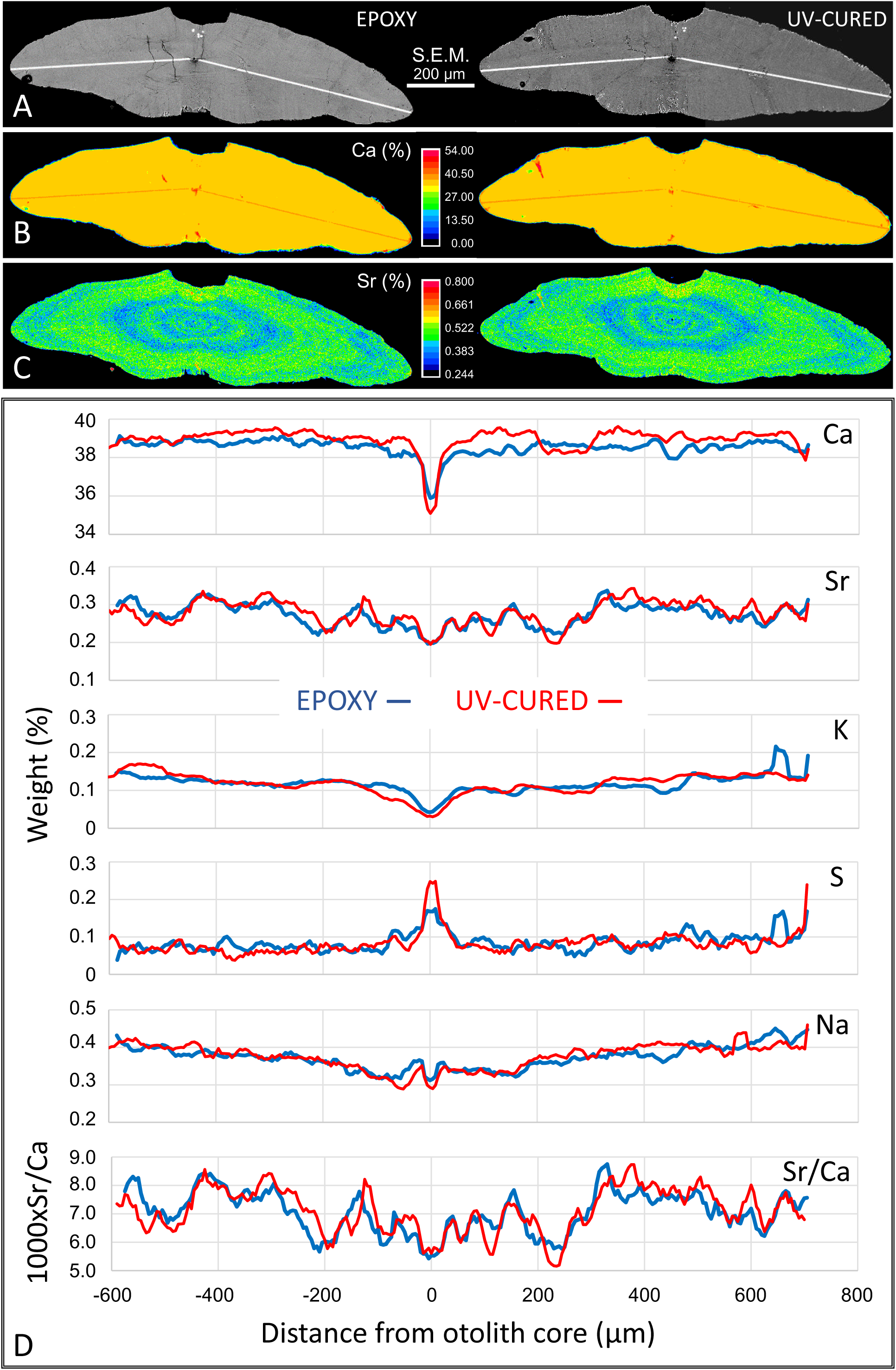
Results of elemental analysis of the contralateral otoliths from one cobaltcap silverside *Hypoatherina tsurugae* that were embedded in UV-cured resin and in Epoxy resin. A) Appearance of the otoliths in SEM view; lines are the marks left by the EPMA Line analysis (spots with 3 μm diameter, 5 μm spacing). B) and C) Results of MAP analysis by EPMA of Calcium and Strontium concentration, respectively. D) Results of EPMA Line analysis of the major elements in cobaltcap otoliths and the calculated Sr/Ca ratios. Both otoliths were aligned with the ventral side to the left for comparison; negative and positive values in the X axis represent the distance from the otolith core to the otolith ventral and dorsal edges, respectively. Results were plotted as the moving average of 5 values.

There are also additional advantages in the use of UV-cured resins. Due to the combination of transparency and hardness, UV-cured resins can be used as a base for otoliths sections that is suitable for direct microscopic observations without a glass slide. This may be relevant also because more often than not otolith cross sections fixed by glues or resins detach from a slide glass during final polishing, observation, or long-term storage. Moreover, this resin base, if molded in an appropriate size and thickness, can also fit a microscope stage or EPMA specimen holder directly (Fig. 2).

## 4. Remaining problems and concerns with UV-curd resins

All of these advantages are not without a price, literally, as UV-cured resins are at present slightly more expensive than the traditional chemically-hardened Epoxy resins. However, prices should theoretically drop in coming years due to the increase in their popularity, diversification of manufacturers, and technological advances in production. A word of caution is also necessary in that little is known about their safety. UV-resins seem to be relatively safe assuming from their growing use (Flexo Magazine (Environotes): <u>http://128.174.142.16/sheets/flexo/uvcuringhealthandsafety.pdf</u>. “Accessed on 4 May 2018”.) but this may simply reflect their novelty and lack of long-term safety studies.

Thus, <u>we strongly emphasize that this report does not constitute an endorsement on their safety. In fact, we strongly recommend the use of laboratory safety and precaution measures (googles, gloves, ventilation, etc) associated with the use of possibly harmful chemicals</u> until more is known on their safety. The same applies to the use of UV-emitting devices for curing these resins. The UV-wavelength required to cure the commercially available UV-resin tested in this study is rather near the spectrum of violet/blue (UVA; 365–400 nm), but even if they were not as dangerous as UVB or UVC, they still have a relatively high energy content and may be associated with the so-called “blue-light hazard”. So before embarking on its use, we suggest that interested readers search for the latest information on the safety of UV-cured resins in public or governmental sites and the respective manufacturer’s MSDS, as well as on the illuminating apparatus/wavelengths used for hardening.

## 5. Concluding remarks

All of these pending safety issues notwithstanding, the authors believe that the use of UV-cured resins may revolutionize otolith specimen preparation practically- and time-wise, and therefore may be extremely useful in situations such as teaching and workshops in which time for otolith embedding is a constraint. Finally, while this report is concerned only with the Acryl Acrylate-based resin, further experimentation with other types of UV-cured resins (e.g. Epoxy- or Vinyl-based resins) may yield even better results than those obtained so far.

## Acknowledgements

The authors acknowledge the TV program “Matsuko no shiranai sekai” (broadcasted 18 Oct 2016) for reporting on the growing use of UV-cured resins for DIY jewelry, which sparked the curiosity of one of the authors (CAS) on the suitability of this medium for otolith embedding. We are also thankful to Ms. Mayumi Otsuki for the careful EPMA analyzes. This work was partially supported by a grant-in aid for scientific research (KAKENHI) to C.A.S. (26241018) from the Japan Society for the Promotion of Science.

## References

Arai, T., Chino, N., 2018. Opportunistic migration and habitat use of the giant mottled eel Anguilla marmorata (Teleostei: Elopomorpha). Sci. Rep. doi:10.1038/s41598-018-24011-z.

Campana, S.E., 1983. Calcium deposition and otolith check formation during periods of stress in coho salmon, Oncorhynchus kisutch. Comp. Biochem. Physiol. A: Mol. Integr. Physiol. 75, 215–220.

Campana, S.E., 1999. Chemistry and composition of fish otolith: pathways, mechanisms and applications. Mar. Ecol. Prog. Ser. 188, 263–297.

Campana, S.E., 2005. Otolith science entering the 21^st^ century. Mar Freshwater Res. 56, 485–495.

Campana S.E., Nielson J.D., 1985. Microstructure of fish otoliths. Can. J. Fish. Aquat. Sci. 42, 1014–1032.

Campana, S.E., Valentin, A.E., MacLellan, S.E., Groot, J.B., 2016. Image-enhanced burnt otoliths, bomb radiocarbon and the growth dynamics of redfish (Sebastes mentella and S. fasciatus) off the eastern coast of Canada. Mar. Freshwater Res. 67, 925–936.

Castellini, D.L., Brown, D., Lajud, N.A., Díaz De Astarloa, J.M., 2017. Juveniles recruitment and daily growth of the southern stock of Mugil liza (Actinopterygii; Fam. Mugilidae): new evidence for the current life-history model. J. Mar. Biol. Assoc. UK. doi: 10.1017/S0025315417001904.

Dilly, P.N., 1976. The structure of some cephalopod statolith. Cell Tissue Res. 175, 147–163.

Fowler, A.J., 1990. Validation of annual growth increments in the otoliths of a small, tropical coral reef fish. Mar. Ecol. Prog. Ser. 64, 25–38.

Grønkjær, P., 2016. Otoliths as individual indicators: a reappraisal of the link between fish physiology and otolith characteristics. Mar. Freshwater Res. 67, 881–888.

Hamilton, S.L., Warner, R.R., 2009. Otolith barium profiles verify the timing of settlement in a coral reef fish. Mar. Ecol. Prog. Ser. 385, 237–244.

Ichii, T., Mugiya, Y., 1983. Comparative aspects of calcium dynamics in calcified tissues in the goldfish Carassius auratus. Nippon Suisan Gakkaishi 49, 1039–1044.

Jones, C.M., 1992. Development and Application of the Otolith Increment Technique. In: Stevenson DK, Campana SE (eds) Otolith Microstructure Examination and Analysis, Canadian Special Publication of Fisheries and Aquatic Sciences 117. Department of Fisheries and Oceans, Canada, pp 1–11.

Kono, N., Takahashi, M., Shima, Y., 2014. Time of formation of incremental and discontinuous zones on sagittal otoliths of larval Japanese Spanish mackerel Scomberomrus niphonius. Nippon Suisan Gakkaishi 80, 21–26.

Morales-Nin B., Bjelland, R.M., Moksness, E., 2005. Otolith microstructure of a hatchery reared European hake (Merluccius merluccius). Fish. Res. 74, 300–305.

Mugiya, Y., 1987. Phase difference between calcification and organic matrix formation in the diurnal growth of otoliths in rainbow trout, Salmo gairdneri. Fish. Bul. 85, 395–401.

Neville V., Rose, G., Rowe, S., Jamieson, R., Piercey, G., 2018. Otolith chemistry and redistributions of Northern cod: evidence of Smith Sound–Bonavista Corridor connectivity. Can. J. Fish. Aquat. Sci. dx.doi.org/10.1139/cjfas-2017-0357.

Radtke, R.L., 1989. Strontium-calcium concentration ratios in fish otoliths as environmental indicators. Comp. Biochem. Physiol. A: Mol. Integr. Physiol. 92, 189–193.

Secor, D.H., Dean, J.M., Laban, E.H., 1992. Otolith Removal and Preparation for Microstructural Examination. In: Stevenson DK, Campana SE (eds) Otolith Microstructure Examination and Analysis, Canadian Special Publication of Fisheries and Aquatic Sciences 117. Department of Fisheries and Oceans, Canada, pp 19–57.

Secor, D.H., Rooker, J.R., 2000. Is otolith strontium a useful scalar of life cycles in estuarine fishes? Fish. Res. 46, 359–371.

Stevenson, D.K., Campana, S.E., 1992. Otolith Microstructure Examination and Analysis, Canadian Special Publication of Fisheries and Aquatic Sciences 117. Department of Fisheries and Oceans, Canada.

Stormer, D.G., Juanes, F., 2016. Effects of temperature and ration on the otolith-to-somatic size relationship in juvenile Chinook salmon (Oncorhynchus tshawytscha): a test of the direct proportionality assumption. Mar. Freshwater Res. 67, 913–924.

Tsukamoto, K., Umezawa, A., Tabeta, O., Mochioka, N., Kajihara, T., 1989. Age and birth date of Anguilla japonica leptocephali collected in western North Pacific in September 1986. Nippon Suisan Gakkaishi 55, 1023–1028.

Watai, M., Ishihara, T., Abe, O., Ohshimo, S., Strüssmann, C.A., 2017. Evaluation of growth-dependent survival during early stages of Pacific bluefin tuna using otolith microstructure analysis. Mar Freshwater Res. 68, 2008–2017.

Watanabe, Y., Kuji, Y., 1991. Verification of daily growth increment formation in saury otoliths by rearing larvae from hatching. Jpn. J. Ichthyol. 38, 1–15.

Zenitani, H., Kono, N., Tsukamoto, Y., Mochioka, N., 2007. Increase of otolith Sr:Ca ratio related to metamorphosis and maturation in anchovy Engraulis japonicus. Fish Sci. 73: 1395–1397.

